# Chronic Antigen Stimulation in Solid Tumors Induces T Cell Exhaustion and Limits Efficacy of T Cell Bispecific Therapies

**DOI:** 10.1101/2025.01.29.635556

**Authors:** Idil Hutter-Karakoc, Eleni Maria Varypataki, Aparna Neelakandhan, Simone Lang, Vesna Kramar, Ahmet Varol, Sasha Simons, Marine Richard, Mudita Pincha, Dario Venetz, Nicole Joller, Christian Münz, Pablo Umana, Christian Klein, Maria Amann

## Abstract

T cell bispecific antibodies (TCBs) have demonstrated promising results in patients with solid tumors. However, the underlying immunological and molecular mechanisms influencing these clinical outcomes require in depth evaluation. T cell exhaustion, a state induced by prolonged antigen exposure, is known to undermine T cell-based immunotherapies, though its specific impact on TCB efficacy remains unclear. In this study, we assessed the effectiveness of TCBs on tumor-specific T cells, focusing on their functional status. Utilizing a fully immunocompetent mouse model with a solid tumor expressing an immunogenic antigen, we showed that tumor-specific T cells acquire an exhausted phenotype and fail to expand under TCB treatment. By employing both mouse and human tumor-specific T cells *in vitro*, our study established that chronically stimulated tumor-specific T cells show impaired response to TCB treatment. The comparison of TCB efficacy in T cell-inflamed tumors with immunogenic antigens versus non-inflamed tumors with low antigen presence in mice revealed TCB success in solid tumors is more reliant on T cell functional fitness than on their abundance before treatment. The data also indicate that solid tumors with elevated levels of both, intratumoral regulatory T cells, and T cells expressing co-inhibitory receptors, show diminished responses to TCB therapy, aligning with similar observations described in hematological cancers. These findings highlight the critical role of T cell exhaustion due to chronic antigen exposure and illustrate that exhausted tumor-specific T cells are likely not the driver population redirected by TCBs for tumor elimination. Our research highlights the importance of maintaining T cell fitness and preventing T cell exhaustion to improve TCB therapy outcomes. This may help better identify patient populations with solid tumors that could benefit from TCB treatments most in clinical settings.

## Introduction

Cancer immunotherapies have transformed the landscape of oncology by harnessing the cytotoxic potential of T cells to specifically target and eliminate cancer cells, delivering remarkable clinical outcomes^1,2^. The prevailing view in the field highlights the presence of tumor-specific cytotoxic T cells as a critical factor for achieving the therapeutic efficacy of immunotherapies ^3,4^. However, a significant challenge remains: tumor-specific cytotoxic T cells frequently enter a state of exhaustion due to chronic antigen stimulation within the tumor microenvironment (TME), diminishing their anti-tumor functionality ^5^ This challenge underscores the need to investigate how their functional profile impacts the efficacy of newer therapeutic classes in cancer immunotherapy.

Over the past two decades, numerous T cell based immunotherapy strategies have been developed, broadly categorized into two subgroups: (i) those relying on cancer elimination via tumor-antigen recognition and (ii) those whose specificity is controlled synthetically^6^. The first group includes early and most widely used immunotherapy approaches such as checkpoint inhibitors (CPis) to reinvigorate repressed T cells, cytokines aiming at the expansion of cytotoxic T cells, cancer vaccines and oncolytic viruses to generate a pool of highly cytotoxic tumor-specific T cells; while the second subgroup consists of advanced antibody engineering techniques like chimeric antigen receptor (CAR) T cells that re-engineer and expand patient’s T cells, and T cell-engaging bispecific antibodies (TCBs) that crosslink cancer cells with the patient’s T cells to facilitate killing independently of their antigen specificity^7,8^.

The infiltration of antigen-specific T cells into tumors is initiated when antigen-presenting cells (APCs) sample the tumor and present tumor antigens to naive T cells in secondary lymphoid tissues. This process primes and activates tumor-antigen-specific T cells, enabling them to infiltrate and eradicate the tumor^9^. Clinical trial findings suggest that tumor T cell infiltration serves as a prognostic marker for cancer immunotherapies^10–13,^ reinforcing the consensus that enhancing the priming and activation of antigen-specific T cells is critical to achieve more effective therapeutic outcomes^14^. However, a vast majority of patients with tumor-specific T cell infiltration still fail to achieve long-term responses to immunotherapies ^2^. This failure can be attributed to T cell exhaustion resulting from persistent T cell receptor (TCR) stimulation due to chronic exposure to tumor antigens^15^, characterized by impaired cytotoxic capacity, reduced cytokine production, and elevated expression of inhibitory receptors^16–18^.

Considering that continuous TCR stimulation might lead to an exhausted phenotype of antigen-specific T cells in the tumor, it is crucial to prioritize alternative strategies such as TCBs that engage not only tumor-specific T cells but also tumor-infiltrating “bystander” cells for anti-tumor immunity^19^. TCBs are bi-specific antibodies with one or more binding moieties recognizing a surface marker on tumor cells, while another binding moiety engages CD3ε on T cells, leading to subsequent T cell activation and tumor cell killing^20–24^. By delivering activation signals through CD3ε, TCBs operate independently of TCR-pMHC recognition, potentially overcoming tumor escape mechanisms related to antigen loss or MHC downregulation^25^. They can also successfully recruit T cells from the periphery into the tumor, thereby increasing T cell infiltration and demonstrating potential for significant efficacy in non-inflamed solid tumors^26^.

Even though TCBs are among one of the most extensively studied bispecific cancer immunotherapies^27^, their efficacy on tumor-specific cytotoxic T cells in solid tumors has not been fully investigated yet. To address this gap, we explored (i) the efficacy of TCBs on tumor-specific T cells, (ii) the influence of the functional profile of these T cells on TCB efficacy, as well as (iii) the role of T cell fitness in therapeutic outcomes from TCBs. Our study demonstrated that tumor-specific T cells do not expand by the TCB treatment *in vivo*. Our findings reveal for the first time that chronically stimulated tumor-specific T cells exhibit an impaired response to TCBs. In line with the findings of the MajesTEC-1 study^28^, we showed that TCBs also promote a more favorable cytotoxic T cell phenotype in solid mouse tumors characterized by lower levels of regulatory T cells (T_regs_) and reduced expression of inhibitory receptors on T cells at baseline.

## Results

### Immunogenic tumor-antigen induces a T cell-inflamed tumor microenvironment with tumor-specific T cells *in vivo*

To study TCB efficacy on tumor-specific T cells in inflamed solid tumors, we first confirmed the endogenous generation of tumor-specific CD8^+^ T cells and their activation status during tumor progression.

To test this, immunocompetent C57BL/6 mice were subcutaneously injected with either OVA^+^ B16 or B16 (OVA B16) melanoma cells. Tumor-infiltrating immune cells were analyzed at specific stages of tumor growth (Figure 1A). The presence of MHC-1-presented OVA antigen resulted in slower tumor progression (Figure 1B). It also led to the generation and infiltration of endogenously formed OVA-specific CD8^+^ T cells which were detected throughout different stages of tumor progression by OVA-Dextramer staining (Figure 1C).

**Figure 1:**
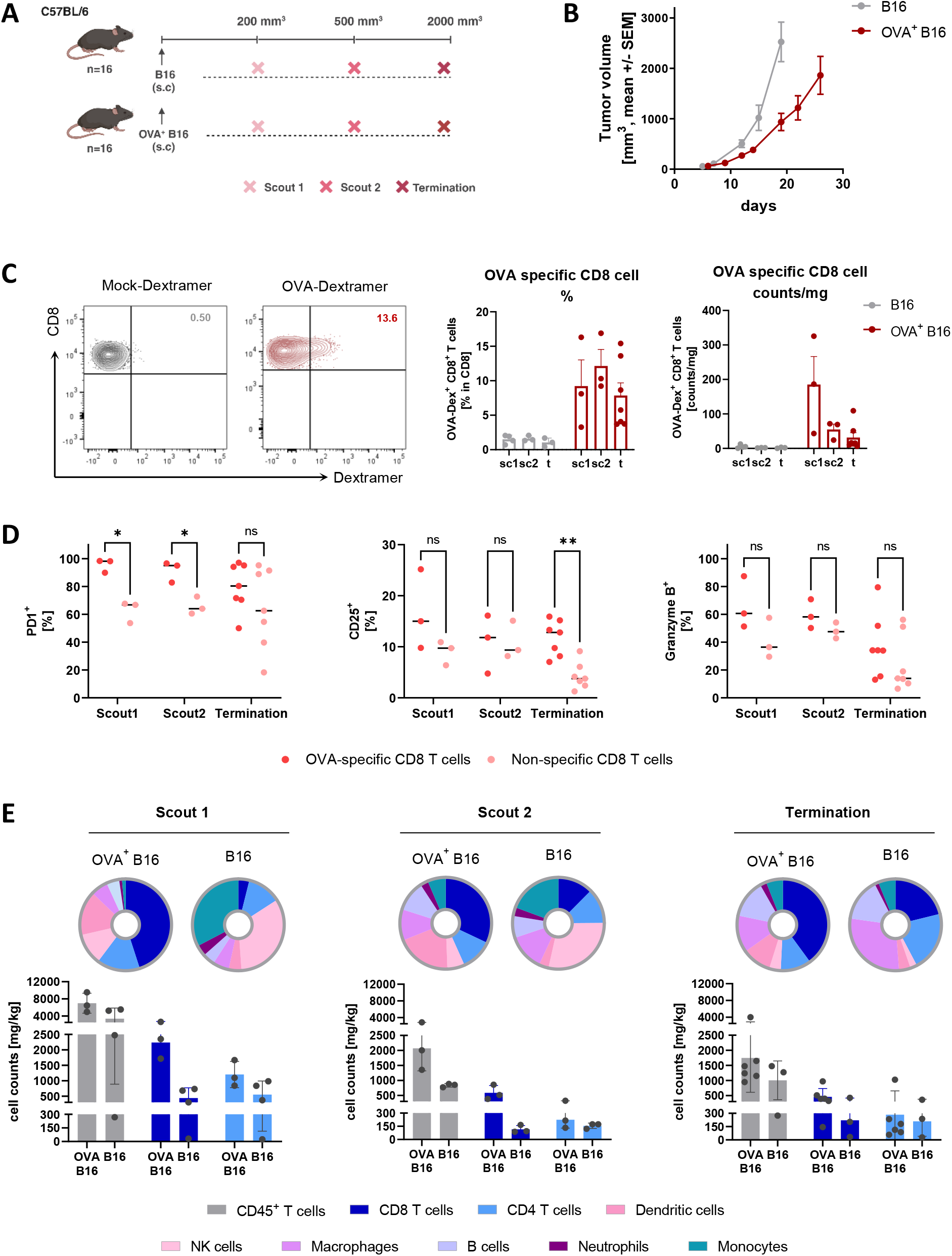
Presence of OVA antigen induces an inflamed tumor phenotype and generates activated antigen-specific T cells in vivo. **(A)** Schematic of the *in vivo* study design. **(B)** Tumor growth curves for subcutaneously injected B16 and OVA^+^ B16 tumors in syngeneic mice (n=16 per group; mean ± SEM). **(C)** Confirmation of endogenous OVA-specific CD8^+^ T cell infiltration of the tumor in the OVA^+^ B16 model on scout 1 (sc1), scout 2 (sc2) and termination (t) timepoints (mean ± SEM). **(D)** Expression levels of PD1, CD25 and Granzyme B in tumor-infiltrating OVA-specific versus non-specific CD8^+^ T cells in the OVA^+^ B16 model (medians indicated as horizontal lines). **(E)** Median abundance (%) of eight different immune cell populations (pie charts, gated on Live CD45+ cells) and quantification of tumor-infiltrating T cells (bar charts) in OVA^+^ B16 and B16 models at scout 1 (200 mm^3^), scout 2 (500 mm^3^), and termination (2000 mm^3^) time points. Each symbol represents a single animal, except in (B) where each dot represents n=8-16 animals. Vertical lines depict mean ± SEM. Statistical comparisons were performed using unpaired t-test with Welch’s correction, *p ≤ 0.05, **p ≤ 0.01, ***p ≤ 0.001.

Next, we assessed the activation and exhaustion status of OVA-specific CD8^+^ T cells. These cells displayed significantly higher levels of activation marker CD25 and inhibitory marker PD-1 compared to non-specific CD8^+^ T cells, suggesting a highly stimulated phenotype (Figure 1D). Anti-gen-stimulated OVA-specific CD8^+^ T also exhibited higher baseline Granzyme B levels (Figure 1D). Furthermore, a decreasing trend in the Granzyme B effector marker was observed in both tumor antigen-specific and non-specific CD8^+^ T cell compartments over the course of tumor progression (Figure 1D). This decline may contribute to explain tumor escape, despite the presence of OVA-specific CD8^+^ T infiltration.

As anticipated, presence of OVA antigen induced not only OVA-specific CD8^+^ T cells infiltration but also T cell-inflamed tumor microenvironment characterized by the infiltration of CD8^+^ and CD4+ T cells (Figure 1E, Supplementary Figure 1A), as well as increased infiltration of eight different detected intratumoral immune cell subsets (Supplementary Figure 1A). Furthermore, OVA^+^ B16 tumors had higher HLA-B expression levels confirmed by histology analysis (Supplementary Figure 2B).

### TCB provides a better cytotoxic T cell phenotype and tumor growth control in non-inflamed tumors *in vivo*

In this study, we aimed to evaluate whether tumors expressing immunogenic antigens, which are recognized by endogenous T cells, exhibit an improved therapeutic response to TCB treatment compared to tumors with little or no tumor antigen expression.

To address this question, murine Tyrp1-TCB (muTyrp1-TCB) was selected as a model TCB, which targets Tyrosinase-related protein 1 (Tyrp1) naturally expressed on the surface of B16 murine melanoma cells (Figure 2A). To compare TCB efficacy in a T cell inflamed tumor with tumor-specific T cell infiltration versus in a non-inflamed tumor model, C57BL/6 syngeneic mice were injected with either OVA^+^ B16 or B16 cells, respectively (Figure 2B).

**Figure 2:**
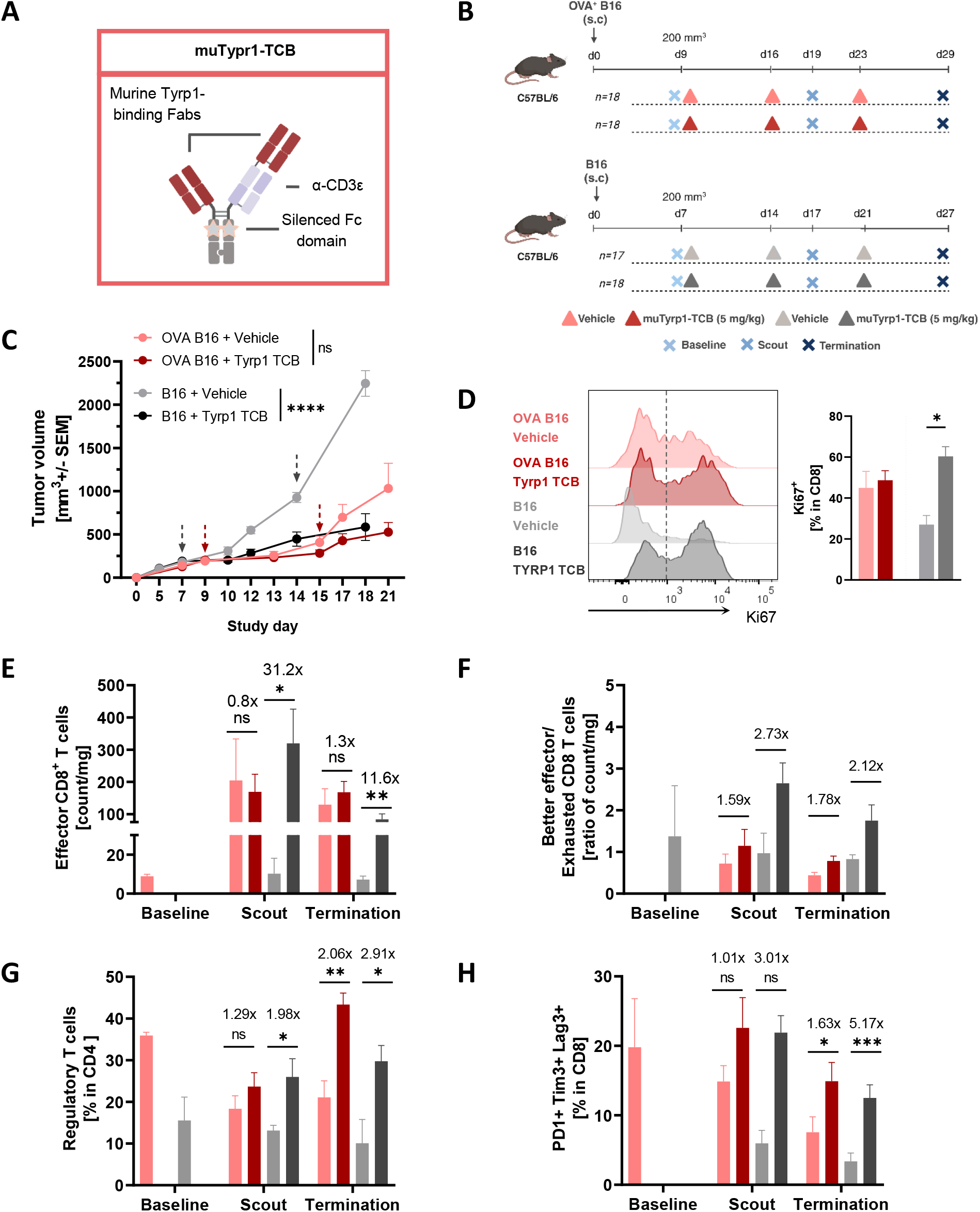
Tyrp1 TCB provides a better cytotoxic T cell phenotype and tumor control in the absence OVA antigen *in vivo*. **(A)** Graphical representation of muTYRP1-TCB design. **(B)** Schematic of the *in vivo* study design. **(C)** Tumor growth of subcutaneously injected OVA^+^ B16 or B16 cells treated with vehicle or muTYRP1-TCB shown till day 21, the last day before B16 vehicle group reached termination criteria. Treatment time points depicted with little arrows for each tumor model: red arrows for Vehicle/muTYRP1-TCB treatment for OVA^+^ B16 model, gray arrows for Vehicle/muTYRP1-TCB treatment for B16 model. (n=8-20 per group; mean ± SEM). **(D)** Ki67^+^ tumor-infiltrating CD8^+^ T cells upon two rounds of muTYRP1-TCB treatment in scout timepoint. **(E)** PD1^+^ TCF1^−^ effector CD8 T cell levels (counts/mg and fold changes) in OVA^+^ B16 and B16 models treated with vehicle or muTYRP1-TCB (mean ± SEM). **(F)** Ratio of Tim3+ Granzyme B^+^ effector CD8^+^ T cells to Tim3+ Granzyme B^-^ exhausted CD8^+^ T cells (counts/mg and fold changes) in the tumor (mean ± SEM). **(G)** Regulatory T cell abundance (% with fold changes) in the TME (mean ± SEM). (H) Co-inhibitory receptor expression levels on tumor-infiltrating CD8^+^ T cells (% with fold changes) upon muTYRP1-TCB treatment over time (mean ± SEM). In (C), each dot represents at least 8 animals. Statistical comparisons were performed using unpaired t-test with Welch’s correction, except for (C) where two-way ANOVA with Sidak test was applied, *p ≤ 0.05, **p ≤ 0.01, ****p ≤ 0.0001.

In the OVA^+^ B16 model, despite the presence of high levels of tumor-infiltrating T cells at baseline (Figure 1E, Supplementary Figure 1A and Supplementary Figure 2A) and high Tyrp1 expression levels over the course of tumor progression (Supplementary Figure 2C), muTyrp1-TCB resulted in only moderate, non-significant tumor growth delay compared to the vehicle control group (Figure 2C and Supplementary Figure 3A). On the other hand, in mice injected with B16 tumors, despite low initial T cell infiltration levels (Figure 1E and Supplementary Figure 1A), muTyrp1-TCB treatment resulted in significant delay in tumor growth and improved survival (Figure 2C). We also analyzed the phenotypic landscape of tumor infiltrating CD8^+^ T cells in both tumor models. In the inflamed OVA^+^ B16 model with infiltrated OVA-specific CD8^+^ T cells, muTyrp1-TCB treatment could not result in a significant increase in proliferative (Ki67^+^) CD8^+^ T cell population (Figure 2D). Also, muTyrp1-TCB treatment did not enhance effector (PD-1+ TCF-1^−^) CD8^+^ T cell infiltration into the tumor compared to vehicle control in this tumor model (Figure 2E). Conversely, in the non-inflamed B16 model, TCB treatment led to significant increase in both proliferative (Ki67^+^) (Figure 2D) and effector (PD-1+ TCF-1^−^) CD8^+^ T cell numbers infiltration at the intermediate (scout) and termination timepoints (Figure 2E). Additionally, mu-Tyrp1-TCB treatment shifted the intratumoral better effector (Granzyme B^+^ Tim-3^+^)/exhausted (Granzyme B Tim-3^+^)CD8^+^T cell ratio in favorof the highly cytotoxic better effector (Granzyme B^+^ Tim-3^+^) cells in the non-inflamed B16 tumors^29^, a shift not observed in inflamed OVA^+^ B16 tumors. (Figure 2F). Notably, non-inflamed B16 tumors exhibited lower infiltration of inhibitory T_reg_ (FoxP3^+^ CD25+) (Figure 2G) and co-inhibitory receptors (PD-1 Tim-3 Lag-3) expressing CD8^+^ T cell (Figure 2H) compared to inflamed OVA^+^ B16 tumors. This observation aligns with previous reports on hematological tumors, indicating that patients with high T_reg_ infiltration and an increased prevalence of exhausted Tcell phenotypes exhibit a diminished response to TCB treatment^28^,^30^.

Taken together, our data demonstrated that TCB treatment can effectively control tumor growth and promote a superior effector phenotype of tumor-infiltrating T cells in a non-inflamed tumor microenvironment. We hypothesized that this superior TCB efficacy in the non-inflamed tumor settings duced the infiltration of T_reg_. and the presence of phenotypically exhausted antigen-specific and non-specific T cells, thereby providing TCB with a more responsive immune population to engage.

### Tumor infiltrating CD8 T cells from inflamed tumor fails to respond to TCB treatment *ex vivo*

*In vivo*, we observed that TCB provided a better cytotoxic T cell phenotype and tumor growth control in the non-inflamed B16 model. To further assess the functional fitness of tumor-infiltrating CD8^+^ T cells (CD8^+^ TILs) *ex vivo*, T cells were isolated at the scout timepoint from both tumor models and co-cultured with B16 cells (Figure 3A). Their ability to respond to muTyrp1-TCB engagement, mediate target cell killing and secrete pro-inflammatory cytokines was evaluated (Figure 3B).

**Figure 3:**
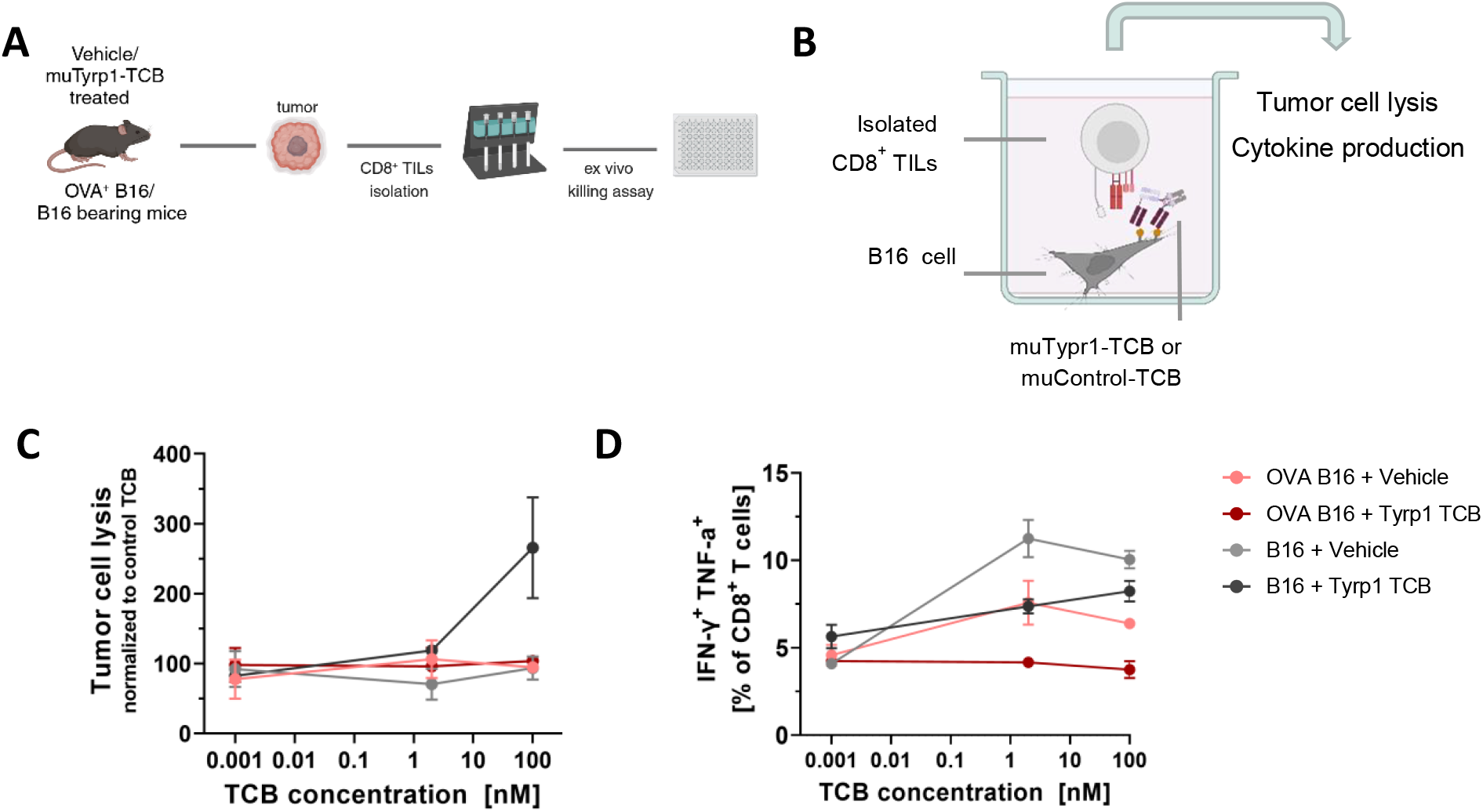
Tumor infiltrating CD8^+^ T cells isolated from OVA^+^ B16 tumor fails to respond to TCB treatment *ex vivo*. **(A)** Graphical representation of *ex vivo* study layout. **(B)** CD8^+^ TILs isolated from the scout timepoint were re-stimulated either with muTyrp1-TCB or with muControl-TCB in the presence of B16 target cells for assessing their tumor cell lysis (24h) and cytokine production capacity (72h). **(C)** B16 cell lysis by CD8^+^ TILs upon muTyrp1-TCB engagement *ex vivo*, normalized to muControl-TCB (mean ± SEM). **(D)** IFN-ɣ and TNF-α production capacity of isolated CD8^+^ TILs upon muTyrp1-TCB engagement with B16 target cells (mean ± SEM). Each dot represents the technical duplicates, which were generated as pooling samples of 4 experimental animals from the same tumor and treatment group. Vertical lines depict mean ± SEM.

Consistent with the *in vivo* findings, CD8^+^ TILs isolated from inflamed OVA^+^ B16 tumors, regardless of the prior *in vivo* treatments, failed to exhibit tumor cell killing *ex vivo* after 24 hours (Figure 3C). In contrast, CD8^+^ TILs isolated from TCB-treated B16 desert tumors demonstrated an enhance tumor cell killing when engaged with muTyrp1-TCB *ex vivo* (Figure 3C).

After an additional 48 hours of incubation, cytokine production was assessed. In alignment with the *in vivo* and *ex vivo* tumor cell lysis results, CD8^+^ TILs isolated from desert B16 tumors, irrespective of their *in vivo* treatment, secreted IFN-γ and TNF-α upon muTyrp1-TCB restimulation (Figure 3D). In contrast, CD8^+^ TILs isolated from vehicle-treated inflamed OVA^+^ B16 tumors exhibited lower IFN-y and TNF-α production, while those from TCB-treated inflamed OVA^+^ B16 tumors completely failed to produce cytokines upon restimulation (Figure 3D).

These data suggest that CD8^+^ TILs from inflamed tumors with persistent antigen presence exhibit a functionally exhausted phenotype, characterized by impaired tumor cell killing and loss of polyfunctionality. These findings are consistent with the results observed in *in vivo* experiments.

### Antigen-specific CD8 T cells do not expand upon TCB treatment *in vivo*

Next, we sought to evaluate the responsiveness of endogenous antigen-specific T cells to TCB treatment *in vivo*.

To investigate this, C57BL/6 syngeneic mice bearing OVA^+^ B16 tumor were treated weekly with 5 mg/kg muTyrp1-TCB or control vehicle. During tumor progression, a set of animals was euthanized and their tumors were analyzed for endogenous OVA-specific T cells (Figure 4A). Strikingly, muTyrp1-TCB did not induce intratumoral expansion of OVA-specific CD8^+^ T cells (Figure 4B and Figure 4C). Over the course of tumor progression these OVA-specific CD8^+^ T cells acquired an effector (PD-1+ TCF-1^−^) phenotype (Figure 4E), followed by subsequent terminal exhaustion (Granzyme R Tim-3^+^), characterized by loss of Granzyme B expression (Figure 4E) ^29^ OVA-specific CD8^+^ T cells also expressed higher levels of co-inhibitory receptor (PD-1 Tim-3 Lag-3) levels associated with T cell exhaustion and dysfunction ^31-33^, compared to non-specific CD8^+^ T cells in the TME (Figure 4F).

**Figure 4:**
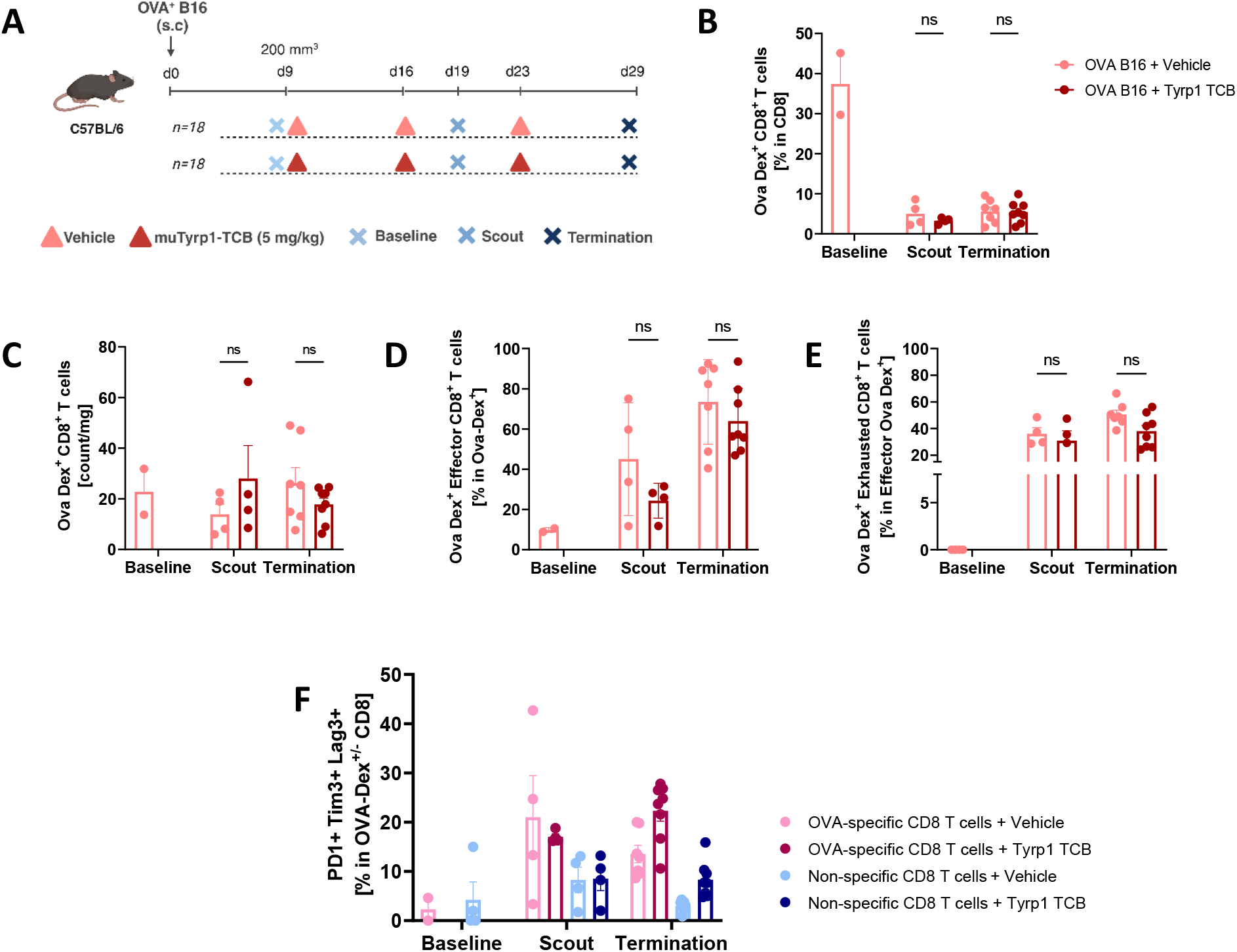
Antigen specific CD8^+^ T cells do not expand upon Tyrp1 TCB treatment *in vivo*. **(A)** Schematic of the *in vivo* study design. Percentages **(B)** and amounts **(C)** of tumor-infiltrating OVA-specific CD8 T cells at baseline, scout, and termination with vehicle or muTyrp1-TCB treatment (mean ± SEM). Infiltration of OVA-specific effector **(D)** and exhausted **(E)** CD8^+^ T cells in the tumor with vehicle or muTyrp1-TCB treatment over time. **(F)** Co-inhibitory receptor expression on OVA antigen-specific vs non-specific CD8^+^ T cells upon muTyrp1-TCB treatment over time. Each dot represents a single animal. Statistical comparisons were performed using unpaired t-test with Welch’s correction, *p ≤ 0.05.

Overall, this data suggest that chronically stimulated anti-gen-specific CD8^+^ T cells in the TME acquire exhausted phenotype and do not expand upon TCB treatment.

### Exhausted mouse CD8 T cells fail to provide anti-tumor immunity upon TCB engagement

Following the *in vivo* observation that chronically stimulated antigen-specific CD8^+^ T cells exhibited exhaustion markers and did not expand upon TCB treatment, we aimed to further assess the impact of functional phenotype of antigen-specific T cells on TCB efficacy *ex vivo*.

For this, we repeatedly exposed OVA-specific murine OT-I CD8^+^ T cells to the OVA_(257-264)_ antigenic peptide *ex vivo* (Figure 5A). As previously reported ^34^, this repetitive antigenic stimulation induces features of exhaustion in the OT-I CD8^+^ T cells (OT-I T_ex_), thereby providing us a model for studying the efficacy of TCBs with exhausted OVA-specific T cells *ex vivo*. As a control, a separate group of OT-I CD8^+^ T cells that were stimulated with the OVA(257-264)Peptide only once, to generate effector OVA-specific T cells (OT-I T_eff_) (Figure 5A). The functional state of generated OT-I T_eff_ and T_ex_ cells was confirmed based on their inhibitory receptor expressions, capacity to proliferate, produce pro-inflammatory cytokines and kill tumor cells. In line with previous findings, OT-I T_eff_ cells showed significantly higher inhibitory receptor expressions (Supplementary Figure 4A-D). In a follow up co-culture with antigen-presenting OVA^+^ B16 tumor cells (Supplementary Figure 4E), OT-I T_ex_ cells exhibited significantly decreased capacity to proliferate (Supplementary Figure 4F), produce pro-inflammatory cytokines IFN-y and TNF-a (Supplementary Figure 4G) and lyse tumor cells (Supplementary Figure 4H) compared to control OT-I T_eff_ cells.

**Figure 5:**
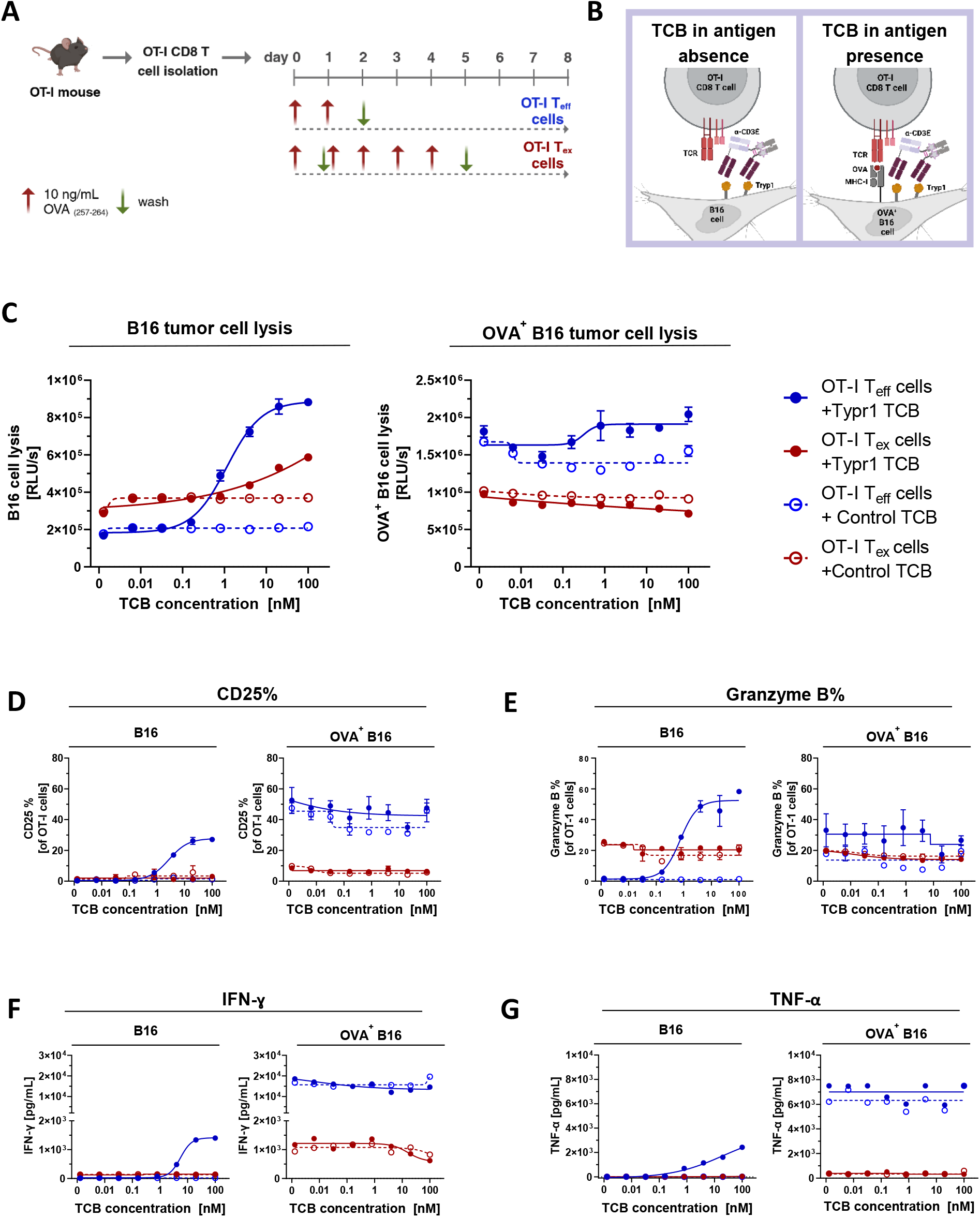
Exhausted OT-I CD8^+^ T cells exhibit impaired response to TCB treatment irrespective of antigen presence. **(A)** Graphical representation of the protocol used to generate activated OT-I T_eff_ and OT-I T_ex_ cells. **(B)** Mechanism of action of muTyrp1-TCB in the presence and absence of OVA antigen on the target B16 tumor cells. **(C)** Lysis of tumor cells by OT-I T_eff_ and T_ex_ cells following 24-hour TCB treatment. **(D)** Expression levels of CD25 in OT-I T_eff_ and T_ex_ cells after 72 hours of TCB engagement with tumor cells. **(E)** Expression levels of Granzyme B in OT-I T_eff_ and T_ex_ cells after 72 hours of TCB engagement with tumor cells. IFN-ɣ **(F)** and TNF-α **(E)** secretion levels in these T cells under the same conditions. Each symbol represents technical duplicates. Data are presented as mean ± SEM. Experiment was performed twice.

After concluding from these data that OT-I T_eff_ cells generated through repetitive antigen stimulation resemble phenotypically exhausted tumor-specific T cells *in vivo*, we wanted to understand how these T cells respond to TCB treatment *ex vivo*. To evaluate the differential response of effector vs exhausted antigen-specific T cells to TCB treatment, OT-I T_eff_ and T_ex_ cells with Tyrp1-expressing B16 tumor cells were cocultured in the presence of either muTyrp1- or a muControl-TCB (Figure 5B). Tumor lysis capacity of these T cells was assessed after 24 hours at increasing concentrations ofTCBs. OT-I T_eff_ cells mediated strong dose dependent tumor cell lysis upon muTyrp1-TCB engagement with less than 1 nM ECso, C_50_ whereas OT-I T_ex_ cells demonstrated reduced killing ability and required more than 7 fold higher TCB concentration (EC_50_=7.6 nM), compared to the muControl-TCB to mediate half maximal lysis (Figure 5C).

We further investigated the impact of chronic antigen exposure on TCR-dependent tumor recognition, as well as the potential modulation of TCB activity by simultaneous engagement with the tumor antigen. For this, OT-I T_eff_ and T_ex_ cells were co-cultured with OVA^+^ B16 target cells, expressing MHC-presented OVA antigen and cell surface receptor Tyrp1 as TCB target (Figure 5B). OT-I T_eff_ cells mediated strong tumor cell lysis upon OVA antigen recognition (Figure 5C), and the addition of Tyrp1-TCB could improve tumor cell killing only slightly (EC_50_=0.1S nM) (Figure 5C). On the contrary, OT-I T_ex_ cells showed reduced tumor cell lysis at baseline; which can be attributed to the observed reduced TCR surface expression levels upon chronic antigen stimulation (Supplementary Figure 4I). They also failed to mediate TCB-dependent killing (EC_50_=17.04 nM) (Figure 5C). These data suggest that regardless of tumor-antigen presence, chronic antigen-stimulated T cells failed to perform TCB directed tumor cell killing.

Following an additional 48 hours of incubation, the expression of the late activation marker CD2S and the directed cytotoxicity marker Granzyme Bon OT-I cells were evaluated.

When co-incubated with B16 tumor cells, OT-I T_eff_ cells displayed a dose-dependent increase in CD2S and Granzyme B (Figure 5D and 5E). In contrast, neither marker expression was induced by TCB treatment in OT-I T_ex_ cells (Figure 5D and 5E). Notably, OT-I T_ex_ cells showed a higher baseline prevalence of Granzyme B (Figure 5E). On the contrary, when OT-I T_eff_ and T_ex_ cells were co-cultured with OVA^+^ B16 tumor cells, the functional capacity seemed to be already saturated by the recognition of the tumor antigen, as no significant differences were detected between the highest (100 nM) and the lowest (0 nM) concentrations of TCB (Figure 5D and 5E).

Cytokine secretion levels were also measured in the supernatants, revealing that OT-I T_eff_ cells produced higher levels of IFN-γ and TNF-α in dose dependent manner, whereas OT-I T_ex_ cells failed to evoke a TCB dependent cytokine response (Figure 5F and 5G). However, it is still worth to note, OT-I T_ex_ cells did exhibit some elevated IFN-γ levels upon antigen recognition on OVA^+^ B16 cells (Figure 5F and 5G). This finding aligns with the considerable levels of intrinsic tumor cell killing observed when co-cultured with OVA^+^ B16 tumor cells (Figure 5C).

As concluded from these experiments, OT-I T_ex_ cells, due to persistent antigen stimulation, failed to mount an effective anti-tumor response upon TCB treatment, regardless of whether their TCRs were engaged with the tumor antigen. In contrast, OT-I T_eff_ cells which were antigen-experienced but not chronically stimulated OT-I T_eff_ cells, demonstrated a strong TCB-mediated anti-tumor immune response, effectively killing tumor cells even in the absence of their cognate antigen. However, there was minimal additional TCB effect observed for tumor cells already presenting their specific antigen.

### Chronic antigen stimulated human MART-1 specific CD8 T cells show impaired efficacy upon TCB treatment

After confirming that chronic antigen-stimulation led to a reduced responsiveness to TCB treatment both *in vivo* and *ex vivo*, we aimed to gain insight into the impact of the functional phenotype of antigen-specific T cells in a human context.

Given the limited availability and fractional presence of human tumor-specific T cells, human MART1-specific T cells were generated through lentiviral transduction with a TCR sequence specific for the human melanoma antigen recognized by T cells 1 (MART-1). Following a previously published protocol^35^, these engineered T cells were exposed to tumor cells presenting the MART-1 antigen under either acute or chronic conditions to derive MART-1 TCR cells with functional phenotypes resembling effector (MART-1 TCR T_eff_) or exhausted (MART-1 TCR T_ex_) states, respectively (Figure 6A).

**Figure 6:**
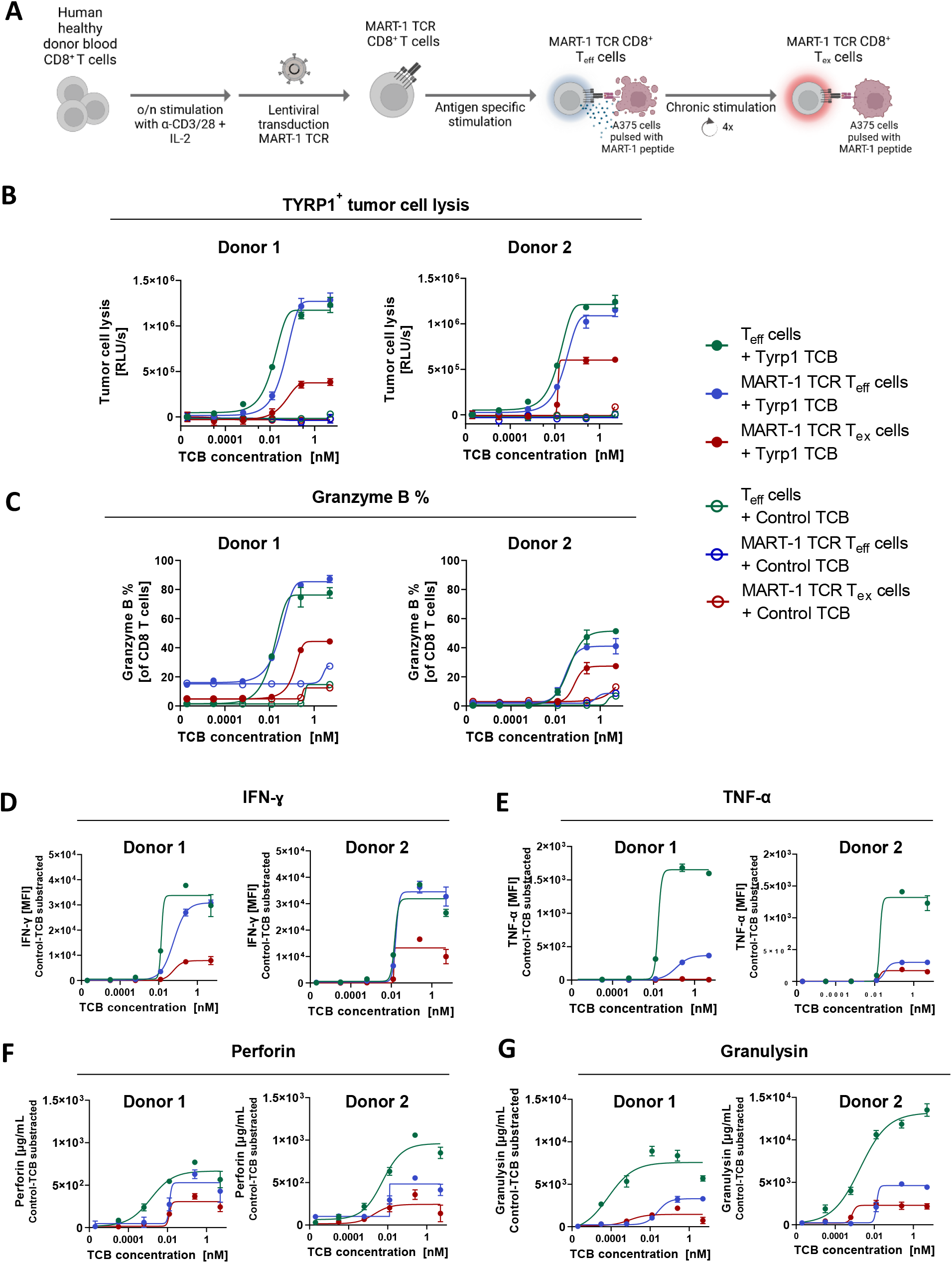
Persistent antigen stimulated human MART1-specific CD8^+^ T cells shows impaired response to TCB treatment in vitro. **(A)** Graphical representation of the protocol used to generate MART-1 TCR CD8 T_eff_ or T_ex_ cells^35^. **(B)** Cytolytic activity of generated MART-1 TCR T_eff_ and T_ex_ cells against CHOK1SV-TYRP1 tumor cells following 48 hours of huTYRP1-TCB treatment. **(C)** Granzyme B expression levels of MART-1 TCR T_eff_ or T_ex_ cells after 48 hours of huTYRP1-TCB engagement with tumor cells. Pro-inflammatory cytokines **(D-E)** and cytotoxic proteins **(F-G)** secretion levels of these T cells under the same conditions. Each symbol represents technical duplicates. Data are presented as mean ± SEM. Experiments were conducted using T cells isolated from two healthy donors.

To investigate the differential efficacy of TCBs on effector versus exhausted human antigen-specific T cells, MART-1 TCR T_eff_ and MART-1 TCR T_ex_ cells were co-cultured with CHOKlSV-Tyrp1 cells expressing the human Tyrp1 receptor. As an additional control, mock TCR-transduced effector T cells (T_eff_) from matching donors were also included. Then, cells were treated with varying concentrations of human Tyrp1 TCB (huTyrp1-TCB) or a human Control-TCB (huControl-TCB). Tumor lysis, cytokine and cytotoxic protein secretion capacity of these T cells were assessed after 48 hours at increasing concentrations of TCBs. In line with *ex vivo* mouse findings, MART-1 TCR T_eff_ cells from two healthy donors mediated a strong dose dependent tumor cell lysis upon huTyrp1-TCB engagement, similar to T_eff_ cells (Figure 6B). In contrast, MART-1 TCR T_ex_ cells from both donors exhibited a reduced tumor cell lysis capacity upon TCB treatment, failing to reach maximal lysis of T_eff_ and MART-1 TCR T_eff_ cells (Figure 6B).

The expression of the directed cytotoxicity marker Granzyme B was evaluated in these cells following hu-Tyrp1-TCB treatment. Both donors’ T_err_ cells and MART-1 TCR T_eff_ cells displayed a dose-dependent increase in Granzyme B expression levels (Figure 6C). Consistent with their observed killing capacity, MART-1 TCR T_ex_ cells demonstrated reduced Granzyme B expression levels upon huTyrp1-TCB engagement (Figure 6C). The secretion levels of cytokines and cytotoxic proteins, including IFN-γ, *TNF-*α, Perforin, and Granulysin, were measured in the supernatants (Figure 6D-6G). Although MART-1 TCR T_eff_ cells were able to kill tumor cells as effectively as non-antigen-experienced T_eff_ cells from the same donors - mock TCR transduced but exposed to CD3/CD28 bead stimulation previously ^-36^they exhibited slightly lower but TCB inducible levels of cytokine and cytotoxic protein secretion (Figure 6D-6F). On the contrary, MART-1 TCR T_ex_ cells showed lower levels of IFN-γ, *TNF-*α, Perforin, and Granulysin secretion levels upon huTyrp1-TCB treatment for both donors.

We also explored additional mechanisms potentially triggered by chronic antigen stimulation which could contribute to the reduced efficacy of TCB in these chronically stimulated antigen-specific T cells. In the *ex vivo* mouse setting, decreased surface TCR expression levels were observed on chronically antigen-stimulated OT-I cells (Supplementary Figure 4). From this observation, we hypothesized that the lower TCR complex levels on chronically stimulated T cells might also lead to decreased CD3 surface expression, which in turn, could potentially result in reduced CD3 availability for TCB engagement in these T cells. To investigate this, TCR and CD3 levels on human MART-1 TCR T cells were quantified during the acute or chronic antigen stimulation, as well as during a subsequent resting period (Supplementary Figure 5). Indeed, although both MART-1 TCR T_eff_ and T_ex_ cells initially exhibited high TCR/CD3 surface expression levels at baseline, they displayed reduced levels following acute or chronic antigen exposure (Supplementary Figure 5). Notably, acutely stimulated MART-1 TCR T_eff_ cells partially recovered their TCR/CD3 surface expression after the resting period, whereas chronically stimulated MART-1 TCR T_ex_ cells maintained lower TCR/CD3 expression levels even after resting (Supplementary Figure 5).

This novel data set confirms that chronic antigen stimulation impairs TCB efficacy in human tumor-specific T cells, highlighting the importance of T cells functional fitness for effective TCB treatments in solid tumors. It also offers insight into a potential explanation for the reduced TCB efficacy on chronically stimulated tumor-specific T cells suggesting it could be due to reduced TCR/CD3 surface expression levels.

## Discussion

TCBs represent a promising and potent class of cancer immunotherapy for solid tumors, offering significant therapeutic benefits to some patients, as demonstrated by the clinical use of the DLL3xCD3 bispecific molecule Tarlatamab^37,38^.TCBs redirect the cytotoxic activity of T cells to cancer cells polyclonally, irrespective of their TCR specificity^19^. Recently, Friedrich et al.^39^ showed that the activation of naive tumor-specific T cells is enhanced when these cells receive TCR-pMHC signaling simultaneously with TCB treatment. However, the field still lacks a comprehensive understanding of the potential influence of T cells’ phenotypic profiles on the therapeutic efficacy of TCBs in solid tumors.

Herein, we explored the efficacy ofTCBs on antigen-specific T cells. Our findings revealed that the functional phenotype of antigen-specific T cells is a key determinant of their response to TCB treatment. Specifically, we observed that TCBs can effectively direct antigen-specific T cells with an effector phenotype towards tumor cell eradication. However, our data illustrate that in solid tumors with chronic antigen stimulation, antigen-specific T cells featuring an exhausted phenotype are unlikely to serve as the primary effector population redirected by TCBs for tumor elimination. Additionally, we showed that antigen-specific T cells were not expanded by TCB treatment as they acquired an exhausted phenotype in the TME. By investigating the TCB efficacy in the presence of tumor-associated antigen, we further showed that antigen presence results in endogenous infiltration of the tumor with inhibitory T_reg_ and exhausted T cells, leading to only a transient response upon TCB treatment in our *in vivo* model. Notably, we also suggest that TCB treatment has the potential to provide a significant antitumor response in non-inflamed tumors, as it can help to enhance cytotoxic effector T cells amounts in the TME.

Clinical results from early cancer immunotherapy trials, particularly with CPis, have established a consensus in the immuno-oncology field that tumors with immunogenic antigen presence, as evidenced by T cell infiltration, are more likely to benefit from these therapies^4,40^ Given that inflamed solid tumors represent only a small fraction of cancer cases^41^we investigated how the efficacy of TCBs differ between inflamed and non-inflamed tumors, and whether pretreatment T cell infiltration is essential for TCB efficacy. Using OVA^+^ B16 and B16 murine melanoma model treated with muTyrp1-TCB as a surrogate TCB allowed us to compare TCB efficacy in T-cell inflamed versus non-inflamed tumor microenvironments side-by-side in a fully immunocompetent mice setting. Despite minimal pre-treatment T cell infiltration, muTyrp1-TCB provided a better cytotoxic T cell phenotype in non-inflamed B16 tumors, characterized by lower T_reg_ and co-inhibitory receptor expressing T cells. This effect was absent in the presence of immunogenic tumor antigen. Our findings, consistent with previous clinical reports on hematological tumors, indicate that the functional fitness of pre-existing T cells, rather than their infiltration levels prior to TCB treatment, is likely the primary determinant of potential TCB efficacy in solid tumors^28,30,39,42^

The impact of infiltrating T cell fitness on TCB efficacy was further confirmed by isolating tumor-infiltrating CD8^+^ T cells from tumors and re-exposing them to the cancer cells upon TCB treatment *ex vivo*. It was evident that CD8^+^ T cells from the inflamed tumor did not exhibit effective anti-tumor responses *ex vivo* either, strongly suggesting a functional impairment of CD8^+^ T cells in the OVA^+^ B16 inflamed model. Our findings highlight that TCBs can be less efficacious in T cell-inflamed, immunogenic tumors, as the high level of immunogenicity can induce tumor-infiltrating T cells to adopt an exhausted phenotype. Taken together, these findings suggest potential benefits of using TCB treatments in non-inflamed solid tumors, where, despite the initially low numbers, T cells exhibiting a functional phenotype could be expanded within the tumor. However, it is also worth mentioning that mouse CD8^+^ T cells may respond differently to the TCB therapy compared to human CD8^+^ T cells. Furthermore, the pathogen-free housing conditions of experimental animals may considerably restrict the diversity and functional phenotype of tumor-infiltrating “bystander” T cells^41,43^potentially impacting the phenotype of infiltrating CD8^+^ T cells and, therefore, the response to the TCB treatment in both models.

In solid tumors, tumor-infiltrating antigen-specific T cells are phenotypically and transcriptionally heterogeneous^41,44^. Therefore, assessing the impact of TCBs on these functionally diverse T cells *in vivo* is of significant importance. To address the question of how TCB treatment impacts antigen-specific T cells in the tumor, we focused on the OVA^+^ B16 tumor model. This model provided a traceable and controlled system to evaluate the efficacy of TCBs on endogenously generated, phenotypically heterogeneous OVA-specific CD8^+^ T cells. Aligning with clinical reporting, in inflamed OVA^+^ B16 model approximately 10 % of the total tumor-infiltrating CD8^+^ T cells were tumor antigen-specific^44^,and these cells exhibited a more exhausted phenotype compared to non-specific bystander T cells. Supported by the finding that OVA-specific T cells are more exhausted than non-specific bystander tumor-infiltrating CD8^+^ T cells, we observed that antigen-specific T cells within the tumors did not increase in numbers following muTyrp1-TCB treatment. Strongly suggesting, these exhausted OVA-specific T cells are not the primary drivers of the anti-tumor response induced by TCB treatment. However, it should be also noted that these data sets were produced using a highly immunogenic OVA antigen potentially amplifying the degree of exhaustion status in the tumor infiltrating antigen-specific CD8^+^ T cell populations, thereby enhancing their diminished response to the TCB treatment.

Antigen-specific T cells represent only a subset of tumor-infiltrating cells, restricting the investigation of TCB efficacy on these cells. To overcome this limitation and study the impact of functional phenotype of antigen-specific T cells on TCB efficacy, we utilized a previously established method to generate functionally distinct effector or exhausted antigen-specific mouse T cells^34^ .Through testing the muTyrp1-TCB on these *ex vivo* generated T cells, we showed that when exhausted, antigen-specific T cells do not respond to the TCB therapy. Our findings demonstrate that unlike effector phenotype OT-I cells, exhausted OT-IT cells can not contribute to anti-tumor immunity upon muTyrp1-TCB therapy. This *ex vivo* finding complements previous reports showing reduced TCB efficacy on exhausted T cells, suggesting that such diminished effectiveness can occur not only in hematological tumors but in solid tumors as well^39, 45^.

To address the limitations associated with using highly immunogenic tumor antigen and mouse models, we next employed human MART-I-specific CD8^+^ T cells, targeting a self melanoma antigen in human tumors^46^. Since antigen-specific T cells of patients constitute a fraction of tumor-infiltrating cells^44^, exhibit varying functional phenotypes^41^, and have different TCR affinities to the same antigen^47^, we generated MART-I TCR-transduced human CD8^+^ T cells from healthy donors. This approach enabled the generation of high of MART-I-specific CD8^+^ T cells, all sharing the same TCR and baseline functional phenotype, with the potential to induce either an effector or exhausted phenotype through single or repetitive antigenic stimulation^35^. Consistent with our observations in the mouse model, chronically antigen-stimulated human MART-I-specific cells exhibited an impaired response to huTyrp1-TCB treatment. As previous studies have shown that TCR-pMHC ligation leads to the ubiquitination and subsequent degradation of the TCR at the protein level^48,49^, and it is also known that, TCR:CD3 complex forms in the endoplasmic reticulum and gets transported to the cell surface as a unit^50^; in our study, we also assessed TCR:CD3 expression levels on MART-I TCR T cells. Our preliminary data suggest that antigen encounter leads to a reduction in CD3-TCR counts on both MART-I TCR T_eff_ and T_ex_ cells. However, while acutely stimulated MART-I TCR T_eff_ cells were able to restore their TCR:CD3 levels after resting in an antigen-free environment, chronically stimulated MART-I TCR T_ex_ cells unable to. Taken together, this dataset highlight that chronic antigen exposure can impair human antigen-specific CD8^+^ T cells by promoting an exhausted functional phenotype and reducing TCR:CD3 surface expression levels, potentially resulting in unresponsiveness to a highly potent TCB treatment.

In this study, we revealed chronic antigen stimulation and subsequent T cell exhaustion as a potential mechanism contributing to resistance to TCB therapy in solid tumors. Our data demonstrate that exhausted antigen-specific T cells, resulting from prolonged exposure to tumor antigens, likely fail to elicit an effective anti-tumor response or expand under TCB therapy. Our findings also indicate that the presence ofimmunogenic antigens can lead to a more exhausted TME, characterized by increased T_reg_ infiltration, accumulation of CD8^+^ T cells expressing co-inhibitory receptors, and reduced TCR:CD3 surface expression which, in turn, can reduce TCB efficacy in inflamed tumors. In light of our findings, we advocate for selecting patients for TCB therapy based on their pre-treatment T cell exhaustion or functional fitness levels, as well as the migratory capacity of their T cells. This approach, rather than relying solely on mutational burden or pre-existing T cell levels, could enhance the therapeutic outcomes of TCBs, particularly in solid tumors, which currently pose significant challenges in the clinics.

## Abbreviations

APC: Antigen presenting cells
CAR: Chimeric antigen receptor
CPI: Checkpoint inhibitor
pMHC: Peptide- Major Histocompatibility Complex
TCB: T cell bispecific antibody
TCR: T cell receptor
TME: Tumor microenvironment
T_reg_: Regulatory T cells
Tyrp1: Tyrosinase-related protein 1

## Acknowledgments

The authors thank all their colleagues from Cancer Immunotherapy, Oncology’. Large Molecule Research and Pharmaceutical Sciences at Roche Pharmaceutical Research and Early Development (pRED) who contributed to this work.

## Author contribution statement

Concept and experimental design: I. Hutter-Karakoc, V. Kramar, A. Neelakandhan, S. Lang, E.M. Varypataki, M. Pincha, M. Amann

Acquisition of data: I. Hutter-Karakoc, A. Neelakandhan Data analysis and interpretation: I. Hutter-Karakoc, E.M. Varypataki, A. Neelakandhan, V. Kramar, A. Varol, S. Simons, M. Richard, C. Klein, C. Münz, N. Joller, M. Amann

Writing, review and/or revision of the manuscript: I. Hutter-Karakoc, V. Kramar, C. Klein, M. Amann Administrative, technical and material support: A. Varol, S. Simons, M. Richard, D. Venetz Study supervision: C. Klein, C. Münz, N. Joller, M. Amann

## Materials and Methods

### Therapeutic Antibodies

All the therapeutic antibodies used in this work were produced internally in Roche.

### Cell lines

OVA^+^ B16Fl0 (Reaction Biology) and B16Fl0 (ATCC) transfected in-house with a plasmid encoding murine fibroblast activation protein (FAP) to generate OVA^+^ B16Fl0 FAP or B16Fl0 cells. The OVA^+^ B16Fl0 FAP cells were cultured in DMEM GlutaMAX (Gibco, 10566016) with 10 % fetal bovine serum (FBS) (Gibco, 26140079), 1.5 µg/mL Puromycin (InvivoGen, ant-pr-1), and 400 µg/mL Hygromycin (Roche, 10566016). B16Fl0 FAP cells were maintained in DMEM GlutaMAX (Gibco, 10566016) with 10 % FBS (Gibco, 26140079) and 0.6 µg/mL Puromycin (InvivoGen, ant-pr-1). CHO-Kl cells (ATCC) and subsequently transfected with a plasmid encoding human Tyrp1 protein 1n-house. The cells were maintained in DMEM GlutaMAX (Gibco, 10566016) supplemented with 10 % FBS (Gibco, 26140079), 6 µg/mL Puromycin (InvivoGen, ant-pr-1).

Lenti-X™ 293T cells (Takara) were maintained in DMEM GlutaMAX (Gibco,10566016) with 10 % FBS (Gibco, 26140079). A375 cells (ATCC) were maintained in DMEM GlutaMAX (Gibco, 10566016) supplemented with 10 % FBS (Gibco, 26140079).

All cell cultures were incubated at 37 °C in a humidified atmosphere containing 5 % CO_2_ and were passaged every 2-3 days.

### Killing assay with *ex vivo* generated OT-I T_eff_ and T_ex_ cells

#### Generation of OT-I T_eff_ and T_ex_ cells

OT-I T_eff_ and T_ex_ cells were generated following the described protocol. ^34^. In brief, OT-I CD8^+^ T cells were isolated from the spleens using negative selection magnetic beads (Miltenyi, 130-104-075) resuspended at 5xl0^5^ cells/mL in mouse T cell media comprising RPMI GlutaMAX (Gibco, 61870036), 10 % FBS (Gibco, 26140079), 1 % Sodium Pyruvate (Gibco, 11360-039), 1 % non-essential amino acids(Gibco, 11140-035), 100 U/mL Penicillin-Streptomycin (Gibco, 15070-063), and 0.05 mM -mercaptoethanol (Gibco, 31350-100). The culture medium was supplemented with murine IL-15 (5 ng/mL, Miltenyi, 210-15), murine IL-7 (5 ng/mL, Miltenyi, 210-07), and daily refreshed 10 ng/mL OVA(257-264) peptide (Anaspec, AS-60193-5). The cells were incubated at 37 °C in a humidified atmosphere containing 5 % CO_2_. After 48 hours, cells were washed three times with PBS (Gibco, 20012027). For OT-I T_eff_cells, they were cultured for 6 more days without OVA(257-264) peptide. For OT-I T_ex_ cells, culture continued for 3 more days with OVA(257-264) peptide added repeatedly.

#### Killing assay set up and luminescence-based readout

OVA^+^ B16Fl0 FAP or B16Fl0 FAP target cells were labeled with PKH-26 Red fluorescent dye (Thermo Fisher, PKH26GL) according to the manufacturer’s instructions. The 40 000 target cells/well (50 µL) were seeded into 96-well U-bottom cell culture plates (TPP, 92697). *Ex vivo* generated OT-I T_eff_ and T_ex_ cells were labeled with 200 nM CellTrace™ CFSE Cell Proliferation dye (Thermo Fisher, 34554) for 10 minutes at 37 °C. The 100 000 CFSE-labeled OT-I T_eff_ or T_ex_ cells were seeded into each corresponding well (50 µL). Pre-diluted murine Tyrp1-TCB or control-TCB were added to the respective wells. Plates were incubated at 37 °C in a humidified atmosphere containing 5 % CO_2_.

After 24 hours, the plates were centrifuged, and 50 µL of the assay supernatant was transferred to a 384-well white plate (Greiner, 655083) before the assay plates were returned back into the incubator. Subsequently, 20 µL of Promega Cytotox-Glo™ reagent (Promega, G9291) was added to each well containing supernatant. The plates were then incubated for 15 minutes at room temperature in the dark. The luminescence signal, corresponding to the relative amount of lysed target cells, was measured using a Tecan SPARK lOM (Tecan).

After an additional 48-hour incubation, plates were centrifuged, and 25 µL of supernatant per well was transferred to new white flat-bottom 384-well plates (Corning, 353988) and stored at -80 °C for later cytokine release assays. Target cells and OT-I cells were for flow cytometry analysis.

#### Flow cytometry readout

The target cells and OT-I cells in the assay plates were transferred to a 96-U bottom plate through rigorous pipetting using a multichannel pipette. Then, cells were washed twice with 200 µL/well PBS (Gibco, 20012027) before staining with 1:1000 diluted LIVE/DEAD™ Fixable Blue Dead Cell Stain (Invitrogen, 123105) at RT for 20 min. After washing the plate once more, cells were stained with a mix of antibodies, to the following surface markers: CD45 (BUV805, BD), CDS (BUV737, BD), PD-1 (BUV395, BD), Lag-3 (BV480, BD), CD25 (PE-CF594, BD), Tim-3 (BV650, BD) for 30 min at 4 °C. Cells were washed with PBS (Gibco, 20012027) and fixed with eBioscience™ Foxp3/ Transcription Factor Fixation Solution (Invitrogen,00-5523-00) (100 µL/well, 1 hour, RT). After the fixation, cells were washed twice with Perm/Wash Buffer (Invitrogen,00-5523-00) and stained with a mix of antibodies diluted in Perm/Wash Buffer (Invitrogen,00-5523-00) to the following intracellular markers: IL-2 (APC/Fire 750, Biolegend), TCF-1 (BV421, BD), Granzyme B (AF700, Biolegend), TOX (PE, Invitrogen), IFN-y (BV786, BD), TNF-a (PE-Cy7, BD) (100 µL/well, 30 min, 4 °C). Cells were washed once with Perm/Wash Buffer (Invitrogen,00-5523-00) and diluted in 100 µL/well PBS (Gibco, 20012027) for acquisition using Symphony A3 flow cytometry (BD).

#### *In vivo* experiments

Female C57BL/6J mice were purchased from Charles River. The experimental study protocol was reviewed and approved by the Veterinary Department of Canton Zurich under the Veterinary License ZHlS0/2020 in agreement with The Swiss Animal Welfare Act. The subcutaneous syngeneic tumor models OVA^+^ B16Fl0 or B16Fl0 were used in this study. In brief, mice were inoculated subcutaneously in the left flank with 2Xl05 OVA^+^B16Fl0 FAP or B16Fl0 FAP cells, mixed 1:1 in Growth Factor-Reduced Matrigel (Corning, 356231) to a volume of 100 µL. Tumor volumes were measured using a caliper, and upon reaching an average volume 200mm3, mice were randomized into treatment groups based on tumor size. Murine Tyrp1-TCB or vehicle control comprising Protein Buffer (Bichsel, 1000366) was administered intraperitoneally at a suboptimal dosage of 5 mg/kg weekly. The animals were euthanized once termination criterion (tumor volume= 2000 mm^3^) was reached.

#### Flow cytometry for *in vivo* samples

Single cell suspension of tumors obtained from OVA^+^ B16Fl0 or B16Fl0 tumor bearing mice were placed in a U-bottom 96-well plate (TPP, 92097) (100 µL/well), and were washed with PBS (Gibco, 20012027). Samples were stained with 1:20 diluted SIINFEKL-loaded MHC Dextramer (APC, Immudex) for 12 min at RT and washed with PBS (Gibco, 20012027). Then, samples were stained with 1:1000 diluted LIVE/DEAD™ Fixable Blue Dead Cell Stain (Invitrogen, L23105) at RT for 20 min in the presence of 1:50 diluted Mouse TruStain FcX Fe Receptor Blocking Solution (101320. Biolegend). Washed samples stained for 30 min at 4 °C in PBS (Gibco, 20012027) containing fluorophore-conjugated antibodies targeting desired surface markers. Before fixation, samples were washed once more with PBS (Gibco, 20012027) and fixed with eBioscience™ Foxp3/Transcription Factor Fixation Solution (Invitrogen, 00-5523-00) (100 µL/well, 1 hour, RT). After the fixation, cells were washed twice with Perm/Wash Buffer (Invitrogen, 00-5523-00) and stained with a mix of antibodies diluted in Perm/Wash Buffer (Invitrogen, 00-5523-00) to the desired intracellular markers (100 µL/well, 1 hour, 4 °C). Samples were washed once with Perm/Wash Buffer (Invitrogen, 00-5523-00) and diluted in 200 µL/well PBS (Gibco, 20012027) for acquisition using Symphony A3 flow cytometry (BD).

For T cell characterization following set of antibodies have been used: CD45 (BUV805, BD), CD8a (BUV737, BD), CD4 (BV510, BD), PDl (BV786, BD), Ox-40 (PE, 119410), CD25 (PE-CF594, BD), Granyzme B (BV421, Biolegend), FoxP3 (AF488, Biolegend) and Ki67 (PE-Cy7, BD). For investigating infiltrating immune cells following set of antibodies have been used: CD8a (BUV737, BD), CD4 (BV510, BD), F4/80 (BV421, Biolegend), Ly6G (BV605, Biolegend), CDllb (FITC, Biolegend), NKp46 (PE, BD), CDllc (PE-CF594, BD), TCRb (PE-Cy5, Biolegend), CD19 (PE-Cy7, Biolegend), CD49b (APC, Biolegend), CD45 (AF700, Biolegend), Ly6c (APC-Cy7, Biolegend).

For investigating muTyrp1-TCB efficacy and its impact on T cells in vivo, following set of antibodies have been used: CD45 (BUV805, BD), CD8a (BUV395, BD), CD4 (BUV496, BD), PDl (BUV737, BD), CD137 (BV510, BD), Tim-3 (BV650, BD), CD127 (BV711, BD), Ox-40 (BV786, BD), Lag-3 (BB515, BD), CD62L (APC/Fire 750, Biolegend), CD25 (PE-CF594, BD), FoxP3 (BV421, BD), TCF-1 (PE, BD), Ki67 (PE-Cy7, BD) and Granyzme B (AF700, Biolegend).

#### Histology

Tumors collected from mice after sacrifice were fixed in 4 % parafolmaldehyde (Thermo Fischer, 160401-AK) overnight. Then, the tumors were embedded in paraffin before immunochemistry stainings. Samples were stained with either a standard H&E protocol using Mayer’s hematoxylin solution (Biosystems, 3870.2500) and eosin 2 % (Biosystems, 84-0023-00), HLA-B (abeam, ab240087) or with an anti-mouse CD3 antibody (Diagnostic Biosystems, RMAB005) following the manufacturer’s instructions. Images were captured with the slide scanner (Olympus, VS200).

#### Ex vivo killing assay

B16Fl0 FAP target cells were labeled with PKH-26 Red fluorescent dye (Thermo Fisher, PKH26GL) according to the manufacturer’s instructions. 5xl04 cells were seeded into a 96 well flat bottom plate (TPP, 92696) in previously described mouse T cell media.

CD8^+^ TILs were isolated from tumor single-cell suspensions using negative selection magnetic beads (Miltenyi, 130-116-478) and debris was removed with a debris removal solution (Miltenyi, 130-109-398). Isolated CDS TILs were counted using a Cedex® HiRes Analyzer (Roche, 05650216001), and lxl0^5^ cells (50 µL/well) were seeded into wells. Pre-diluted TCBs were added at 50 µL/well. Plates were incubated at 37 °C with 5 % CO_2_.

After 24 hours, the plates were centrifuged, and 50 µL of the supernatant was collected. Then, 20 µL of Promega Cytotox-Glo™ reagent (Promega, G9291) was added to the supernatant. After a 15-minute incubation in the dark at room temperature, luminescence was measured using a Tecan SPARK lOM (TECAN), indicating the relative amount oflysed target cells. Following an additional 48 hours of incubation, target cells and TILs were stained for flow cytometry analysis.

### Killing assay with MART-1 T_eff_ and T_ex_ cells

#### CD8 T cell isolation from buffy coat

Buffy coats were ordered from Blutspende Zurich. PBMCs were isolated from the buffy coats using Leucosep tubes (Greiner, 227290) filled with 15 mL Histopaque density gradient medium (Sigma-Aldrich, 10771) according to manufacturer’s recommendations. CD8^+^ T cell isolation was performed by negative selection using the CD8^+^ T cell isolation kit (Miltenyi, 130-096-495). Cells were cultured in base human T cell media comprising advanced RPMI (Gibco, 12633-012), 10 % FBS (Sigma, F4135-500ML), 1 % Glutamax (Gibco, 35050-038). The media was supplemented with 25 ng/mL IL-7 (Miltenyi, 130-095-364) and 50 ng/mL IL-15 (Miltenyi, 130-095-766).

#### Preparation of virus-like particles (VLP)

Lipofectamine LTX™ (Invitrogen, 15338500) based transfection of ∼60 % confluent Lenti-X™ 293T cells (Takara, 632180) was performed with MART-I TCR (DMF5) as well as packaging vectors pCAG-VSVG and psPAX2 at a 2:1:2 molar ratio. As control for every experiment, mock virus-like particles (VLPs) using only the packaging vectors, but no transfer vector, were produced. After 48 hours, the supernatant was collected, centrifuged for 10 minutes at 500 x g to remove remaining cells and then concentrated 10-fold (Lenti-X-Concentrator, Takara, 631231) by centrifugation and resuspension according to the manufacturer’s protocol.

#### Generation of MART-1 TCR T cells

Isolated CD8^+^ T cells were seeded at lxl06 cells/mL in G-Rex® 24-well cell culture plates (Wilson Wolf, 80192M) and activated using Immunocult T cell Activator cocktail (Stem-cell Technologies, 10990) for 16-24 hours.

VLPs (150-300 µl) were added together with 8 µg/ml Polybrene (Sigma Aldrich) and Lentiboost P (1:100) (Sirion Biotech, SB-P-LV-101-12) to the activated T cells. Transduction efficacy was assessed by flow cytometry earliest 72 hours after transduction. Assays were usually performed between day 7-15 post transduction.

MART-I TCR T_eff_ and T_ex_ cells were generated following a previously published protocol3^5^• A375 cells (25 000/well) were seeded in 96-well plates and pulsed with 2 µM MART-I peptide (MEL, SP0009). After 4-16 hours, cells were washed with PBS and human T cell media (100 µL/ well) was added. MART-I TCR T cells (100 000)were then co-cultured with the peptide-pulsed A375 cells, supplemented with 50 U/mL Proleukin. For T_ex_ cells, T cells were re-challenged with peptide-pulsed A375 cells over 9 days with 4 restimulation rounds. For T_eff_ cells, T cells were challenged once for 72 hours. Cells were rested overnight and normalized to cell counts before assay use.

#### Killing assay set up and luminescence-based readout

CHOKlSV-TYRP-1 cells were harvested, washed, and seeded into 384-well V-bottom (Thermo Scientific, 4309) plates at 40 000 cells/well in 40 µL of base human T cell media. MART-I T cells, normalized to counts, were added to the CHO-Kl TYRP-1 cells at the same density. Pre-diluted human Tyrp1-TCB or control-TCB were added, reaching a final volume of 100 µL per well and the desired concentration. The plates were incubated at 37 °C in a humidified atmosphere with 5 % CO_2_.

After 48 hours, the plates were centrifuged, and 20 µL of the assay supernatant was transferred to white flat-bottom 384-well plates (Corning, 353988). For the assessment of cytotoxic activity, 10 µL of Promega Cytotox-Glo™ reagent (Promega, G9291) was added to each well. The plates were incubated for 15 minutes at room temperature in the dark. Luminescence, indicative of the relative amount oflysed target cells, was measured using a Tecan SPARK lOM (TECAN).

For the cytokine release assay, 30 µL of the assay supernatant was transferred to new white flat-bottom 384-well plates (Corning, 353988) and stored at -80 °C. Subsequently, MART-1 cells were stained for flow cytometry analysis.

#### Flow cytometry for killing assay

Cells in the assay plates were washed twice with 70 µL/well PBS (Gibco, 20012027) before staining with 1:1000 diluted LIVE/DEAD™ Fixable Blue Dead Cell Stain (Invitrogen, L23105) at RT for 20 min. After washing the plate once more, cells were stained with a mix of antibodies, to the following surface markers: CD45 (BUV805, BD), CD25 (BB515, BD), Lag-3 (PerCP-Cy5, Invitrogen), CD127 (PE-Cy5, Biolegend), CD107a (BV605, Biolegend), Tim-3 (BV711, BD), CD69 (BV750, BD), PDl (BV786, BD) (40 µL/well, 30 min, 4 °C).

Cells were washed with PBS (Gibco, 20012027) and fixed with Cytofix/Cytoperm™ Fixation/Permeabilization solution (BD, 554714) (30 µL/well, overnight, 4 °C). After the fixation, cells were washed twice with Perm/Wash Buffer (BD,554714) and stained with a mix of antibodies diluted in Perm/Wash Buffer (BD,554714) to the following intracellular markers: Granzyme B (RB780, BD), TCF-1 (APC, CST), CD3 (AF700, Biolegend), Ki67 (BV421, Biolegend). Cells were washed once with Perm/Wash Buffer (BD, 554714) and diluted in 100 µL/well PBS (Gibco, 20012027) for acquisition using Symphony A3 flow cytometry (BD).

#### Cytokine measurement

Cytokines secreted from T cells during the killing assays were analyzed from the frozen assay media. Cytokines were analyzed using either BD™ Cytometric Bead Array (CBA) kits (BD), Bio-Plex Pro Mouse Cytokine 8-plex Assay (BIO-RAD, M60000007A) or LEGENDplex™ Human CD8/NK Panel (13-plex) w/ VbP V02 kits according to manufacturer’s recommendation.

#### Receptor quantification

Receptors on MART-1 T cells were quantified using Quantum™ Simply Cellular® (Bangs Laboratories, 815B) kit according to manufacturer’s recommendation.

#### Data Analysis

Flow Cytometry data was analyzed using FlowJo V.10.10. GraphPad Prism V.9.5 was used to generate the graphs for statistical analysis. The statistical tests used are indicated in the figure legends.

#### Illustrations

Illustrations shown in this work were created using Biorender.com

