## Supplementary Figures for "Chronic Antigen Stimulation in Solid Tumors Induces T Cell Exhaustion and Limits Efficacy of T Cell Bispecific Therapies"

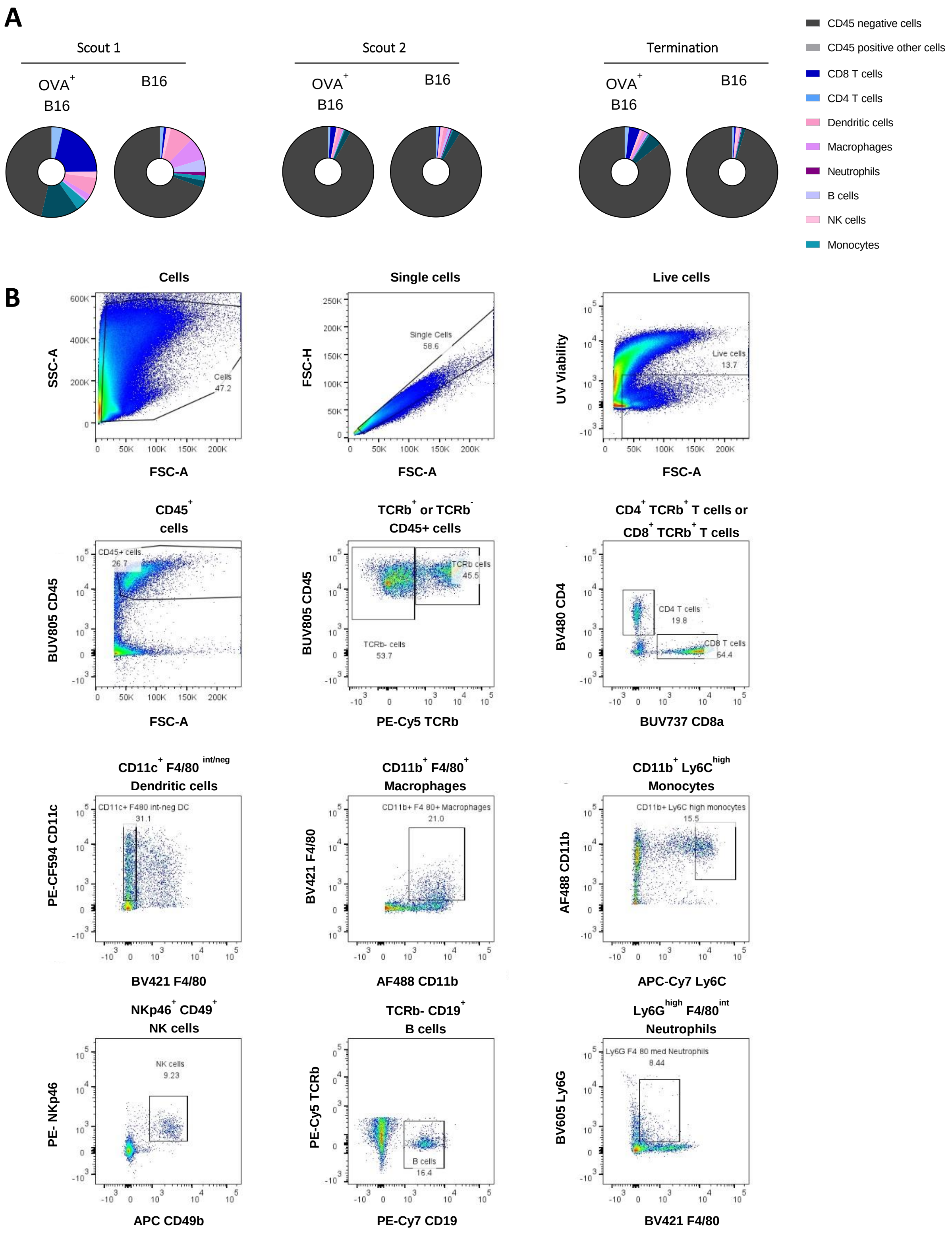


**Supplementary Figure 1: Immunogenic antigen presence in OVA B16 in vivo model led to T cell inflamed tumor microenvironment.**

**(A)** Median abundance (%) of eight different immune cell populations within total live cells. **(B)** Gating strategy of eight different immune cell populations in tumor.

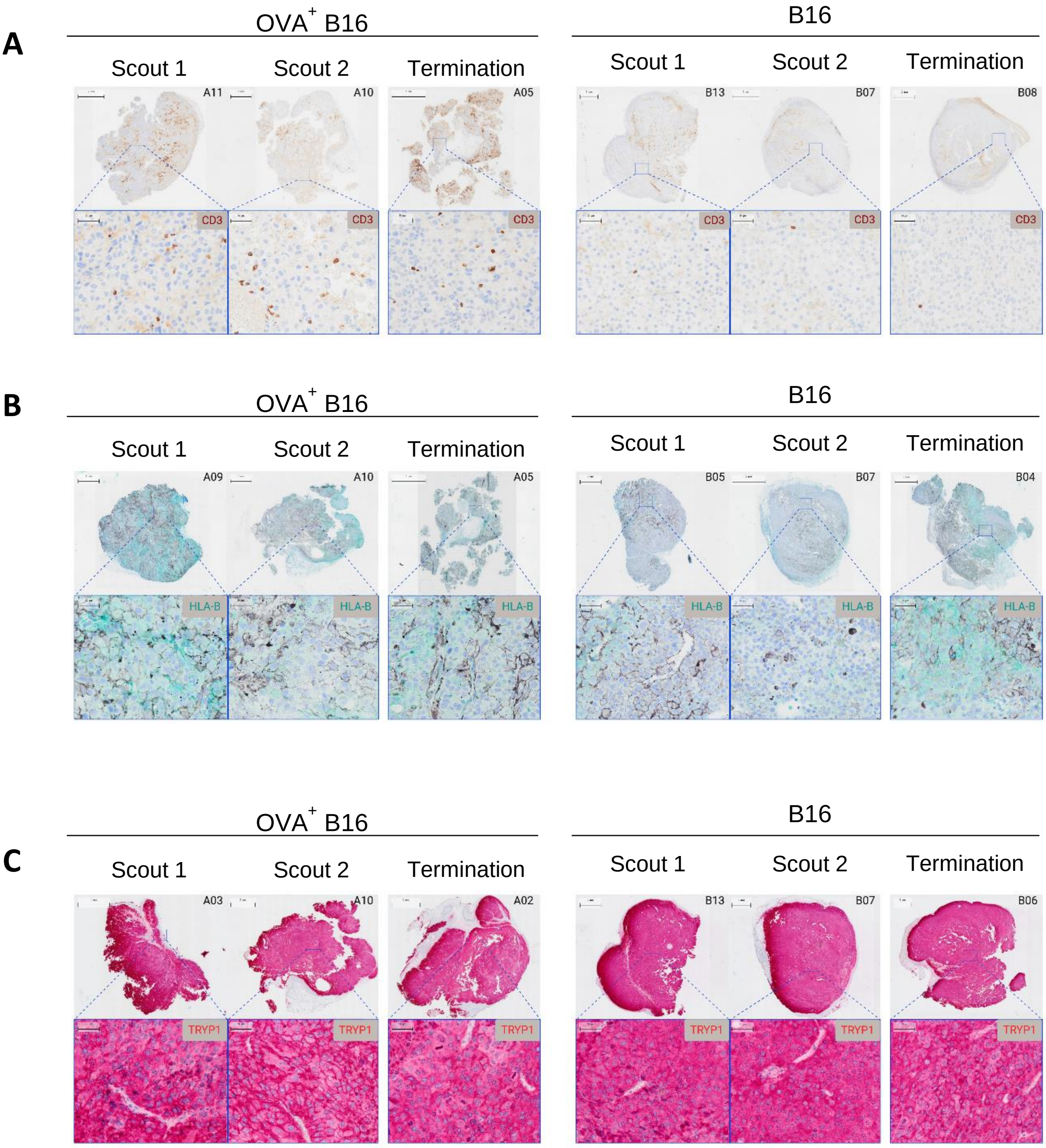


**Supplementary Figure 2: Confirmation of T cell infiltration, HLA-B and Tyrp1 expressions over the course of in vivo study for OVA$^{+}$ B16 and B16 FAP models.**

**(A)** CD3$^{+}$ T cell infiltration levels (brown). **(B)** murine HLA-B expression levels (blue). **(C)** Murine Tyrp1 expression levels (pink).

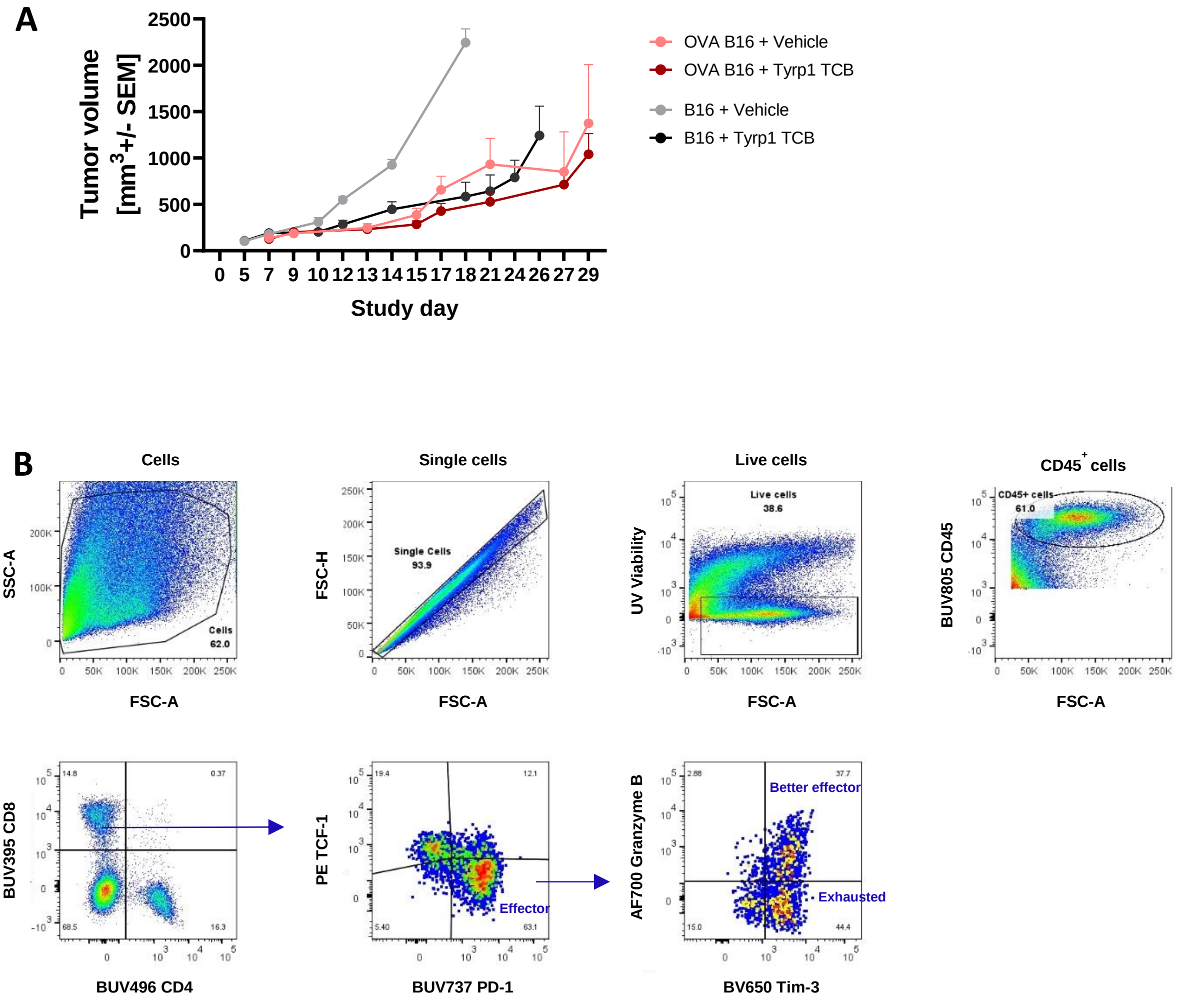


**Supplementary Figure 3: Tumor growth control and T cell gating strategy in B16 and OVA B16 models.**

**(A)** Tumor growth control on both B16 and OVA B16 models provided by muTYRP1-TCB over the course of in vivo study. **(B)** Gating strategy of tumor infiltrating $CD8^+$ T cells.

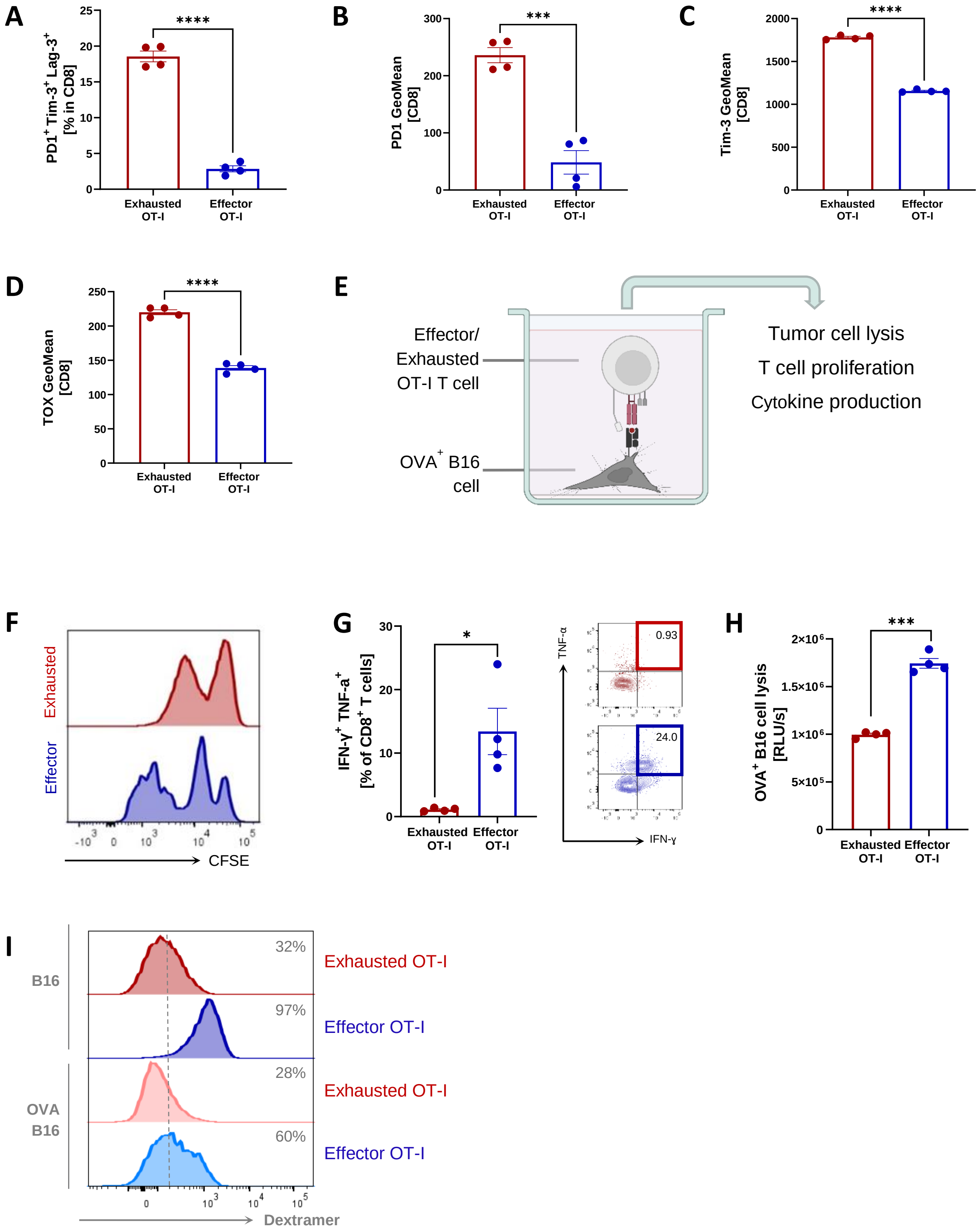


**Supplementary Figure 4: Chronic antigens stimulated OT-I $T_{ex}$ cells express high levels of inhibitory receptors and fail to produce cytokines**

**(A)** Expression of inhibitory receptors on OT-I CD8+ T cells 3 days after chronic antigen stimulation (day 8). GeoMean signal of PD1 **(B)**, Tim3 **(C)** and TOX **(D)** on OT-I CD8+ T cells 3 days after chronic peptide stimulation (day 8). **(E)** Experimental setup for co-incubation of OT-I T cells with antigen-presenting OVA+ B16 cells to evaluate T cell proliferation (72h), cytokine production (72h) and tumor cell lysis (24h). Proliferation **(F)**, cytokine production **(G)** and tumor cell lysis capacity **(H)** of OT-I $T_{eff}$ and $T_{ex}$ cells upon co-incubation with OVA+ B16 cells. **(I)** TCR expression levels of OT-I T cells after co-culturing with B16 or OVA+ B16 cells, detected by OVA-Dextramer staining. Each symbol represents a technical replicate. Data are presented as mean ± SEM. Statistical comparisons were performed using an unpaired t-test with Welch's correction, *$p \leq 0.05$, ***$p \leq 0.001$, ****$p \leq 0.0001$.

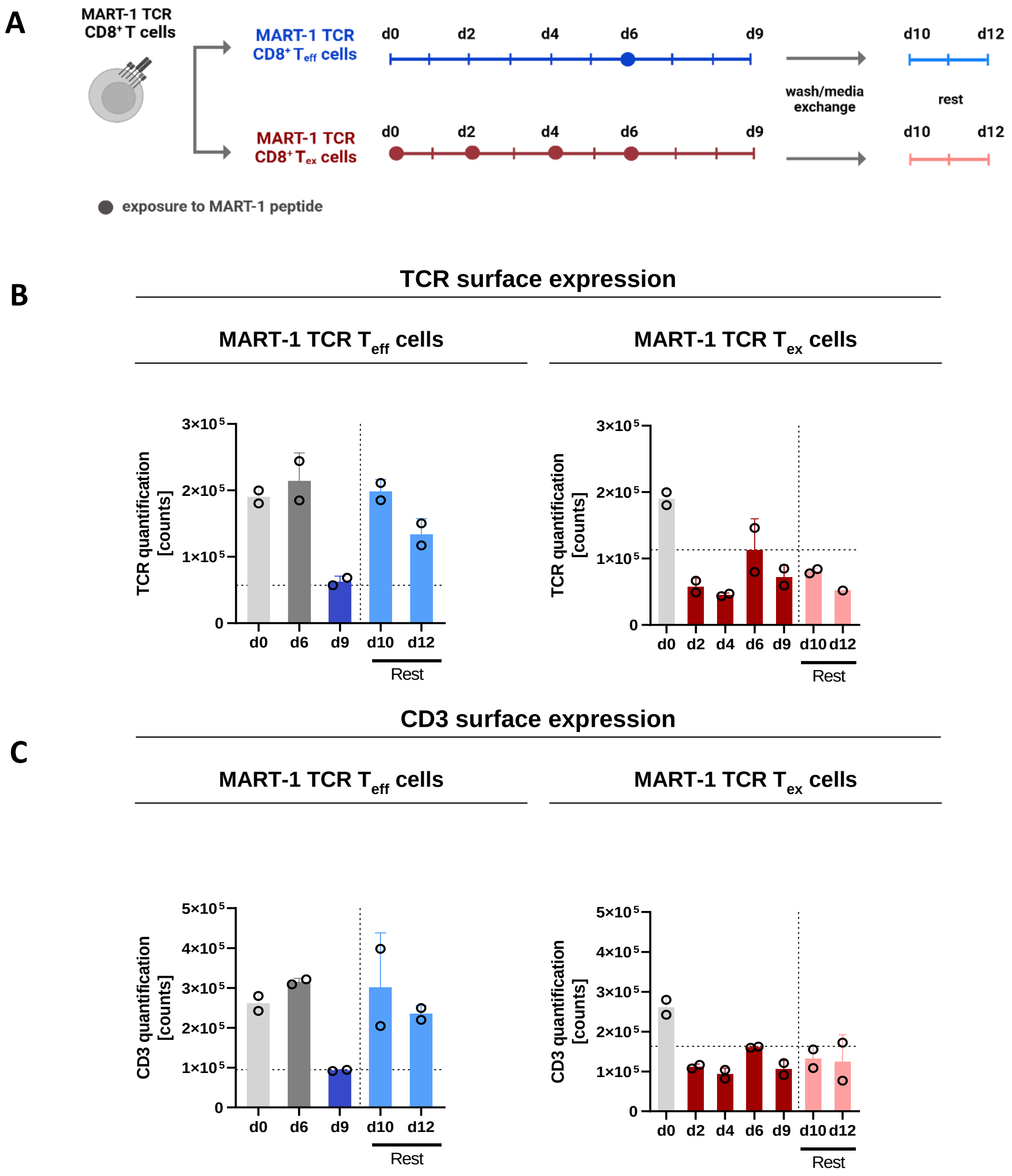


**Supplementary Figure 5: Chronic antigen stimulated human MART1-specific CD8$^{+}$ T cells shows reduced CD3 and TCR surface expression levels *in vitro***

**(A)** Graphical representation of the time points for quantifying CD3 and TCR surface expression. Quantitative analysis of TCR **(B)** and CD3 **(C)** surface expression on MART-1 TCR $T_{eff}$ and $T_{ex}$ cells at baseline (day 0), during (days 2-9), and post (days 10-12) MART-1 peptide stimulation. Each data point represents the average TCR or CD3 TCR expression from a minimum 3000 cells of one donor. Experiments were conducted using T cells isolated from two healthy donors.
